# Altered volume-regulated anion channel activity contributes to depression and anxiety-related molecular and behavioural phenotypes in zebrafish

**DOI:** 10.64898/2026.07.22.740018

**Authors:** Athira Ajith, Durai Shalu, Rudrakant Sharma, Gautam Mohapatra, Amal Kanti Bera

## Abstract

Major depressive disorder (MDD) is a leading cause of global morbidity and mortality. Unfortunately, a substantial proportion of patients do not respond adequately to currently available therapies, highlighting the need for new therapeutic targets. Here, we investigated the role of volume-regulated anion channel (VRAC) in depression-related phenotypes using zebrafish. Disruption of VRAC function, either by pharmacological inhibition or morpholino-mediated knockdown of *lrrc8aa*, the zebrafish ortholog of mammalian *LRRC8A*, the obligatory subunit required for VRAC function, induced anxiety- and depression-like behaviours in zebrafish larvae and altered the expression of genes associated with affective disorders. Transcriptomic analysis of *lrrc8aa*-deficient larvae revealed dysregulation of pathways involved in neuronal signalling and cellular stress responses. Conversely, pharmacological activation of VRAC with zinc pyrithione (ZPT) improved behavioural abnormalities and partially restored altered gene expression. In adult zebrafish subjected to chronic unpredictable stress, ZPT produced antidepressant- and anxiolytic-like effects comparable to those of imipramine and normalized elevated monoamine oxidase (*mao*) expression. Together, these findings indicate that reduced VRAC function contributes to depression- and anxiety-related behavioural and molecular phenotypes and identify VRAC as a potential target for the development of novel antidepressant therapies.

**Graphical abstract:** 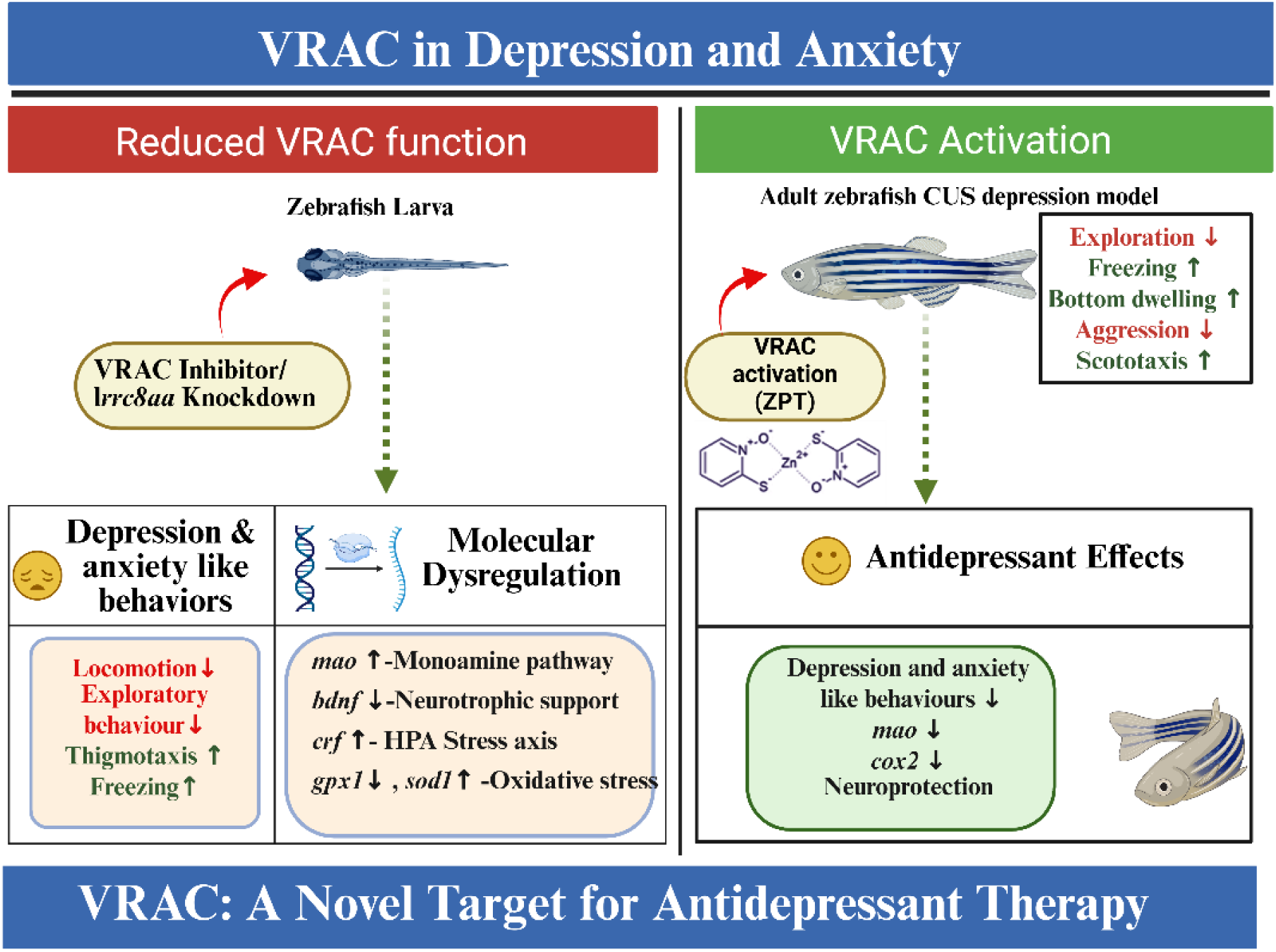

## Introduction

Major depressive disorder (MDD) is a leading cause of disability worldwide and affects millions of people across all ages and sexes (Cui et al., 2024; McCarron et al., 2021). Although several classes of antidepressants are available, many patients either remain symptomatic despite treatment or relapse after discontinuation. Long-term use of these drugs is also associated with significant side effects (Cartwright et al., 2016). Together, these limitations highlight the need to identify new therapeutic targets and develop more effective treatments.

MDD arises from complex interactions among genetic, hormonal, and environmental factors (Yuan et al., 2023). Classical hypotheses attribute depression to reduced monoamine signaling, particularly serotonin, norepinephrine, and dopamine, or to dysregulation of the hypothalamic– pituitary–adrenal axis with excess cortisol production (Malhi and Mann, 2018). In parallel, growing evidence implicates altered neuronal excitability, impaired synaptic plasticity, and chronic neuroinflammation (Duman et al., 2016; Hassamal, 2023; Rominto et al., 2025). Ion channels play a central role in regulating neuronal activity, and their dysfunction has been linked to several neuropsychiatric conditions (Imbrici et al., 2013). Among cation channels, variants in genes encoding calcium channels, potassium channels, and the P2X7 receptor have been associated with mood disorders (Huang et al., 2024). For example, polymorphisms in KCNK2, which encodes the TREK1 potassium channel, have been linked to MDD, and KCNQ2/3 potassium channel openers show antidepressant potential (Congiu et al., 2015; Costi et al., 2022; Zhang et al., 2024). Notably, ketamine, a rapid-acting antidepressant, exerts its effects by blocking NMDA-type glutamate receptors (Krystal et al., 2024, 2023). In contrast, anion channels have received far less attention in this context, despite their critical role in neuronal excitability and signaling. This gap represents an important and largely unexplored avenue for antidepressant drug development.

Volume-regulated anion channels (VRACs) are swelling-activated chloride channels that mediate regulatory volume decrease and contribute to cellular homeostasis (Qiu et al., 2014; Voss et al., 2014). Functional VRACs are heterohexameric complexes composed of the essential LRRC8A subunit together with one or more LRRC8B–E subunits (Thöne et al., 2025). In the brain, VRACs are widely expressed and regulate osmotic balance, neuronal excitability, and the release of neuroactive molecules such as glutamate, aspartate, and taurine (Kimelberg et al., 1990; Mongin, 2016). Astrocytic VRACs support neuron–glia communication under physiological conditions but can drive excitotoxic signaling under pathological states (Yang et al., 2019). Pharmacological inhibition of VRAC reduces glutamate release and attenuates stroke-induced brain injury, and astrocyte-specific LRRC8A knockout mice are protected against ischemic stroke (Alibrahim et al., 2013; Yang et al., 2019; Zhang et al., 2008). VRAC has also been implicated in synaptic plasticity, learning, and memory. Astrocyte-specific depletion of VRAC decreases presynaptic glutamate release and impairs hippocampus-dependent learning and memory (Yang et al., 2019). Although VRAC dysfunction has been associated with epilepsy, ischemia, and traumatic brain injury, its role in major depressive disorder remains largely unexplored (Yang et al., 2025).

To address this gap, we used zebrafish (*Danio rerio*) as a tractable model to examine whether VRAC contributes to depression- and anxiety related behaviours. Zebrafish are a well-established vertebrate model for neuropsychiatric research, sharing substantial genetic similarity with humans and possessing conserved neurochemical pathways. They display well-characterised behavioural endpoints relevant to depression and anxiety, respond to clinically used antidepressants, and express functional VRAC throughout development (Costa et al., 2023; Siddiqui et al., 2025; Yamada et al., 2016; Yang et al., 2025).

Here, we show that both pharmacological inhibition and genetic suppression of VRAC induce depression-and anxiety-related phenotypes in zebrafish larvae. We further demonstrate that pharmacological activation of VRAC alleviates behavioural and molecular abnormalities in larval and adult zebrafish models. These findings suggest that reduced VRAC function contributes to affective dysregulation and identify VRAC as a potential therapeutic target for mood disorders.

## Materials and Methods

### Animals

All experimental procedures were approved by the Institutional Animal Ethics Committee (Approval No. IITM/IAEC-5/AKB-01/2025). Wild-type *Danio rerio* were maintained in a recirculating aquaculture system under controlled conditions (27–28 °C, pH 7.4 ± 1, 14:10 h light/dark cycle). Breeding was conducted at a 2:1 male-to-female ratio, with males and females separated overnight and spawning induced at the onset of light. Embryos were collected within 15 min post spawning and maintained in E3 medium. Following behavioural experiments, larvae and adult fish were euthanized by ice water immersion. Larval heads and pooled adult brains were collected and stored at −80 °C for subsequent molecular and histological analyses.

### DCPIB dose response and Embryotoxicity Assessment

Embryos at 4 dpf were exposed to increasing concentrations of 4-(2-butyl-6,7-dichlor-2-cyclopentyl-indan-1-on-5-yl) oxybutyric acid (DCPIB) (5-30 μM) for 24 h at 27-28 °C. Vehicle-treated embryos served as controls. LD₅₀ was estimated using nonlinear regression analysis.

### Morpholino Oligonucleotide Injection

Translation-blocking antisense morpholino oligonucleotides (MOs) targeting the ATG start site of lrrc8aa (NM_001029949.1; 5′-ACCGCAGCTCAGTGATGGGAATCAT-3′) were obtained from Gene Tools. A Scrambled MO (5’-CCTCTTACCTCAGTTACAATTTATA-3’) was used as a control (Gene Tools). Morpholinos were diluted in Danieau buffer (58 mM NaCl, 0.7 mM KCl, 0.4 mM MgSO₄, and 0.6 mM Ca(NO₃)₂ in distilled water; pH 7.6–7.8) supplemented with 0.05% phenol red. Approximately 1 nl of MO solution was injected into fertilized embryos at the one-cell stage.

### Electrophysiology

Zebrafish embryos (24–48 hpf) were dissociated into single cells using protease and trypsin– EDTA. The dissociated cells were maintained at room temperature and used for electrophysiological recordings within 3 h of preparation. Whole-cell VRAC currents were recorded using an Axopatch 200B patch-clamp amplifier, as described previously (Yamada et al., 2016). Cells were held at −60 mV in isotonic extracellular solution (ECS) and subsequently superfused with hypotonic solution for approximately 5 min to achieve full activation of VRAC. Currents were elicited by applying voltage steps from −100 to +100 mV in 10 mV increments. Recordings were obtained first in hypotonic ECS and then in hypotonic ECS supplemented with ZPT (30 μM)

### Larval Behavioural Assays

Larvae were exposed to pharmacological treatments prior to behavioural testing. At 5 dpf, individual larvae were transferred to either 6-well or 96-well plates (one larva per well) and allowed to habituate before open-field assessment. Touch evoked escape responses were elicited by gently stimulating the tail with a fine needle. Locomotor activity and anxiety-like behaviour were recorded for 10 min using the ZebraBox system (ViewPoint Life Sciences, France). Recordings in 6-well plates were used to quantify exploratory rate and thigmotaxis, taking advantage of the larger arena, and were analyzed using ToxTrac software. Recordings in 96-well plates were used to assess total distance travelled and freezing behaviour using ViewPoint software. The thigmotaxis index was calculated as the ratio of distance travelled near the periphery to the total distance travelled. The exploratory rate was defined as the percentage of area explored [(area explored / total area) × 100]. Freezing duration was defined as the total time spent immobile (with swimming speed below 2 mm/s for at least 1 s, using the default ViewPoint ZebraLab settings) during the recording period.

### Chronic Unpredictable Stress (CUS) Paradigm and Drug Treatment

Adult male zebrafish were randomly assigned to experimental groups. The CUS protocol consisted of exposure to two randomly scheduled stressors per day for 15 days, including restraint, social isolation, overcrowding, tank changes, thermal stress, chasing, net elevation, water change, and dorsal body exposure. Details of the CUS protocol are provided in the Supplementary Material (Table 1). From day 5 of the CUS paradigm, zebrafish were treated with ZPT (1 mg/L), imipramine (1 mg/L), or zinc chloride (1 mg/L) via immersion for 1 h twice daily for 10 consecutive days following stress exposure.

### Adult Behavioural Assessment

Behavioural testing was conducted between 09:00 and 17:00 h following a 30 min habituation period. All sessions were recorded and analyzed using EthoVision XT 17 (Noldus, Netherlands). The Novel Tank Test (NTT) was used to assess anxiety-like behaviour and locomotor activity. After a 5 min acclimation period, individual fish were introduced into a tank divided into equal top and bottom zones, and behaviour was recorded for 10 min. Vertical exploration, freezing (complete absence of movement), and locomotor activity were quantified. Anxiety- and depressive phenotypes were inferred from increased bottom-zone preference, higher frequency of entries into the lower zone, and prolonged freezing.

Aggression and social interaction were assessed using the mirror-induced aggression test. Time spent near the mirror and the frequency of mirror-directed attacks were quantified, with reduced interaction interpreted as indicative of increased anxiety- or depressive states. The scototaxis (light/dark preference) test was used to further evaluate anxiety related behaviour. Fish were allowed to freely explore light and dark compartments for 10 min, and increased preference for the dark zone was interpreted as elevated anxiety- or depression-like behaviour.

### RT-qPCR analysis

cDNA was synthesised from total RNA using the GoScript™ Reverse Transcription System (Promega, Madison, WI) and gene expression was analysed by RT-qPCR with SsoAdvanced Universal SYBR Green Supermix (Bio-Rad, Hercules, CA, USA). Relative gene expression was calculated using the 2^-ΔΔCT^ method with β-actin as the reference gene. Primer sequences are provided in Supplementary Material (Table 2).

### Transcriptomic analysis

RNA-seq reads were aligned to the *Danio rerio* reference genome (GRCz11; GCA_000002035.4) obtained from Ensembl. Differential gene expression between control and *lrrc8aa* morphant larvae was analyzed using DESeq2 (Love et al., 2015). Genes with a log_2_ fold change (log_2_FC) ≥ 1 and a false discovery rate (FDR) < 0.05 (Benjamini–Hochberg correction) were considered differentially expressed; genes with log_2_FC ≥ 1 were classified as upregulated, and those with log2FC ≤ −1 as downregulated. Gene Ontology (GO) and KEGG pathway enrichment analyses were performed using SRplot, with the full *Danio rerio* transcriptome as the background and an FDR < 0.05 as the significance threshold.

### Western Blotting

Protein lysates from 24 hpf embryos were prepared in RIPA buffer (pH 8.0) supplemented with PMSF and protease inhibitors (1:100; Sigma-Aldrich). Protein concentrations were determined using the Bradford assay. Equal amounts of protein were separated by SDS–PAGE and transferred onto PVDF membranes. Membranes were blocked for 1 h in Tris-buffered saline containing 0.1% Tween-20 and 5% BSA, followed by overnight incubation at 4 °C with rabbit anti-LRRC8A (Sigma, SAB1412855; 1:1000) and mouse anti-β-actin (Sigma-Aldrich, A5316; 1:5000). After incubation with HRP-conjugated secondary antibodies, signals were detected using Clarity ECL substrate (Bio-Rad) on a ChemiDoc XRS system. Band intensities were quantified using ImageJ.

### Histology

Cresyl violet staining was used to assess morphological changes and structural damage in zebrafish brain tissue (telencephalon). Paraffin-embedded brain sections were first deparaffinized with xylene. Then it was subjected to a series of ethanol washes. Sections were then stained with 0.1% cresyl violet solution, rinsed with distilled water, and cleared again in xylene. Finally, the sections were mounted with DPX, cover slipped, and imaged using an upright microscope.

### Statistical Analysis

Data are presented as mean ± SEM. Statistical analyses were performed using GraphPad Prism. Data distribution was assessed using the Shapiro–Wilk test. For normally distributed data, comparisons between two groups were performed using unpaired Student’s *t*-tests, while multiple group comparisons were analyzed using one-way ANOVA followed by Tukey’s post hoc test. For non-normally distributed data, two-group comparisons were conducted using the Mann–Whitney U test, and multiple group comparisons were analyzed using the Kruskal – Wallis test followed by Dunn’s post hoc test. Statistical significance was defined as *P* < 0.05 (*\*P < 0.05; **P < 0.01; ***P < 0.001; ****P < 0.0001, ns-non significant).* Unless otherwise stated, *n* represents individual animals and *N* denotes independent biological replicates.

## Results

### Pharmacological inhibition of VRAC induces depression- and anxiety-like behaviours in zebrafish larvae

To investigate the role of VRAC in larval behaviour, we pharmacologically inhibited VRAC using DCPIB. Although DCPIB has been reported to exert off target effects, it remains the most widely used and comparatively selective inhibitor among all currently available VRAC blockers (Decher et al., 2001; Jentsch et al., 2016; Yang et al., 2019). Zebrafish larvae were exposed to increasing concentrations of DCPIB for 24 h to determine LD₅₀ and developmental abnormalities. The LD₅₀ was found to be approximately 18.57 µM (Fig. 1A). DCPIB concentrations above 12 µM induced severe developmental toxicity in zebrafish larvae, characterized by gross morphological abnormalities including spinal curvature, body axis defects, tail malformations, and pericardial/yolk sac oedema (“monstrous phenotypes”) (Fig. 1B and 1C). Therefore, 10 µM, which produced minimal morphological abnormalities, was selected for behavioural experiments. Four days post-fertilization (dpf) larvae were treated with 10 µM DCPIB for 24 h, after which locomotor behaviour was recorded using an automated tracking system (Zebrabox) and analyzed with Toxtrac or ViewPoint software. Ethanol-treated larvae served as vehicle controls. DCPIB-treated larvae displayed normal touch evoked escape responses, indicating preserved motor capacity (Supplementary Video 1). However, quantitative behavioural analysis over a 10-min period revealed a significant reduction in total distance travelled and exploratory activity compared with controls (Fig. 1D, 1F and 1G). Further, DCPIB-treated larvae exhibited a marked increase in freezing duration (Fig. 1E) and thigmotaxis (time spent near the arena boundary) (Fig. 1H), behaviours commonly associated with depression- and anxiety related states in zebrafish.

**Figure 1.**
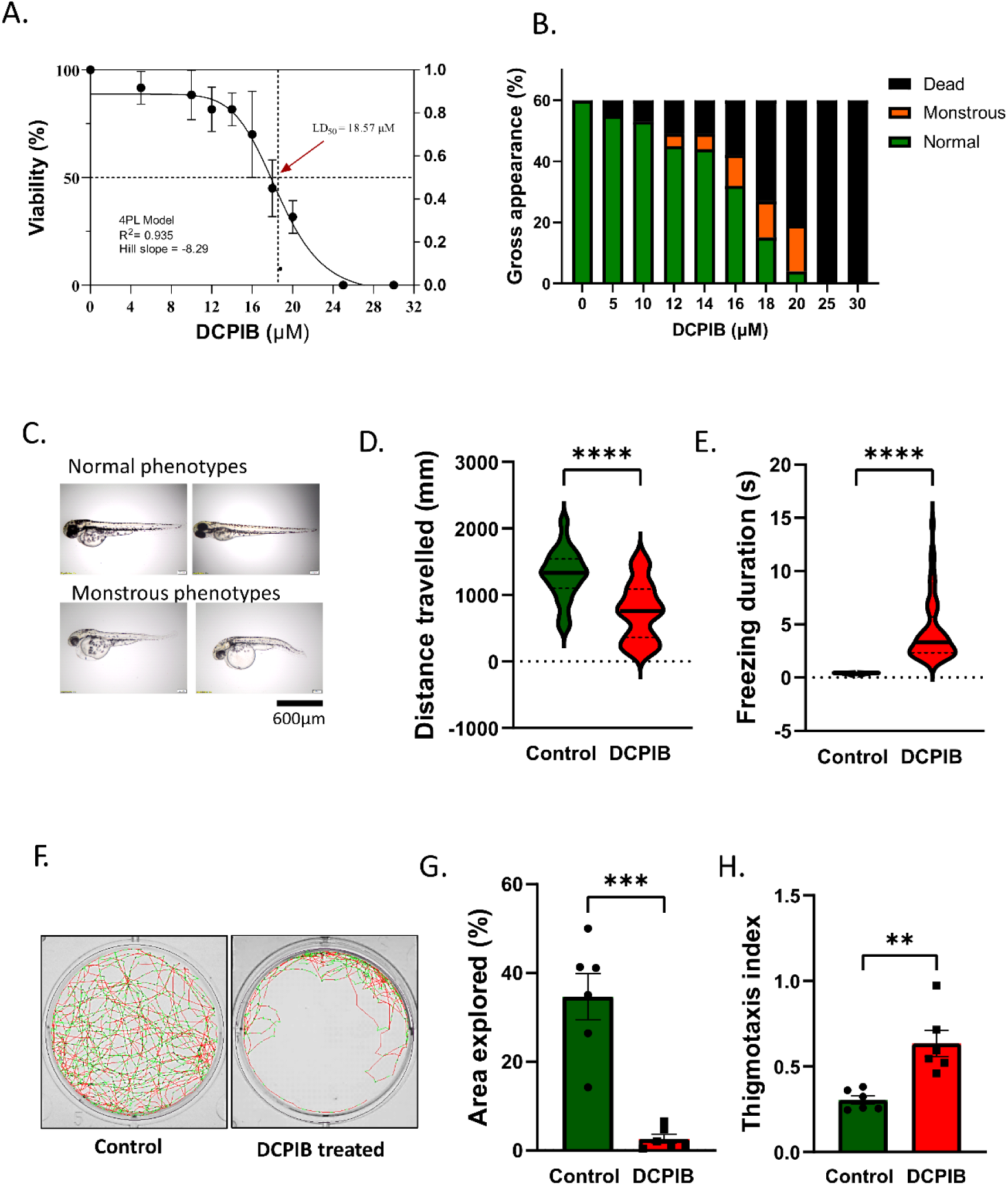
Pharmacological inhibition of VRAC induces behavioural alterations in zebrafish larvae. (A) Survival of 4 dpf larvae following 24 h DCPIB exposure (5–30 μM). (B) Distribution of phenotypic outcomes (dead, monstrous, or normal larvae) across DCPIB concentrations. (C) Representative images of normal and DCPIB-treated larvae showing developmental abnormalities. Scale bar, 600 μm. (D, E) Quantification of total distance travelled and freezing duration from 96-well locomotor recordings (n = 48 larvae/group). (F) Representative locomotor trajectories of control and DCPIB-treated larvae in the open-field assay. (G, H) Quantification of exploration rate and thigmotaxis index from 6-well recordings (n = 6 larvae/group). Data are presented as mean ± s.e.m.; statistical analyses were performed using Mann–Whitney U test (D, E) or unpaired Student’s t-test (G, H).

### Morpholino mediated *lrrc8aa* depletion induces depression- and anxiety-like behaviours

To further validate the role of VRAC in larval behaviours, we reduced LRRC8A protein levels using a translation-blocking morpholino targeting *lrrc8aa*, the zebrafish ortholog of human *LRRC8A*. Because LRRC8A is an obligatory subunit of VRAC, its reduction decreases the number of functional VRAC channels. Morpholinos were injected at the one-cell stage, and embryos were allowed to develop into larvae. Morpholino-injected larvae exhibited varying degrees of deformities. In severe cases, embryos at 24 to 48 hpf showed gross developmental abnormalities, including body axis defects, impaired tail extension, pericardial oedema, and disrupted somite formation (Fig. 2B). Approximately 80% of larvae injected with 0.5 mmol/L *lrrc8aa* morpholino developed normally, whereas 5% died and 15% exhibited severe abnormal phenotypes. Both mortality and malformation rates were higher than in the control morpholino group (Fig. 2A). Knockdown efficiency at the protein level was assessed 24 to 30 hours post-injection by western blotting. Larvae injected with 0.5 mmol/L *lrrc8aa* morpholino showed approximately 50% reduction in LRRC8A protein levels (Fig. 2C).

**Figure 2.**
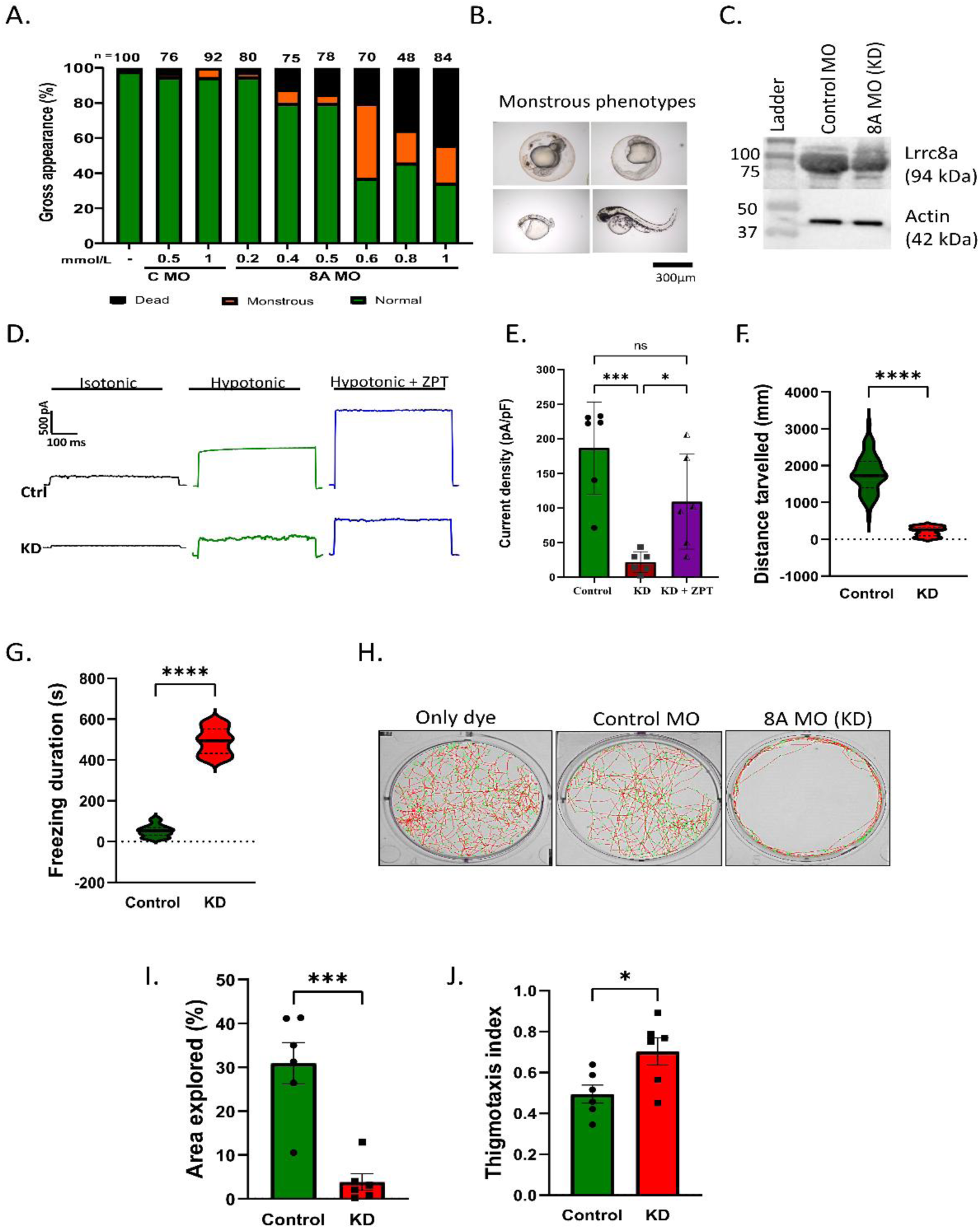
Morpholino-mediated depletion of *lrrc8aa* induces depression- and anxiety-like behaviours in zebrafish larvae. (A) Phenotypic distribution of embryos injected with control (C) or *lrrc8aa* morpholino (MO) at indicated concentrations, classified as dead, monstrous, or normal. Numbers above bars indicate total embryos analyzed. (B) Representative images of developmental abnormalities following *lrrc8aa* knockdown. Scale bar, 300 μm. (C) Western blot showing LRRC8A protein levels in control and *lrrc8aa* MO-injected larvae. Actin served as a loading control. (D) Representative VRAC current traces recorded from control and *lrrc8aa* morphants in hypotonic solution with ZPT. (E) Mean VRAC current density at +100 mV in control, knockdown, and knockdown + ZPT groups (n = 6 cells). (F–G) Behavioural analysis in 96-well plates showing (F) total distance travelled and (G) freezing duration (n = 48 larvae/group). (H) Representative locomotor trajectories of dye-injected, control MO-, and *lrrc8aa* MO-injected larvae. (I–J) Quantification of (I) exploration rate and (J) thigmotaxis index from 6-well recordings (n = 6 larvae/group). Data are presented as mean ± s.e.m. Statistical analyses are described in Methods. KD, knockdown.

For functional validation, VRAC currents were measured using whole-cell patch clamp in zebrafish embryonic cells injected with control or *lrrc8aa* morpholino. As shown in Fig. 2D and E, *lrrc8aa* knockdown cells exhibited significantly reduced VRAC currents upon exposure to hypotonic solution compared to control cells. Treatment with the VRAC activator ZPT (30 μM) restored VRAC activity in knockdown cells, although not to control levels.

Larvae injected with *lrrc8aa* morpholino were used for subsequent behavioural analyses. Both control larvae and apparently normal *lrrc8aa*-depleted larvae displayed intact touch-evoked responses (Supplementary Video 2). However, VRAC deficient larvae at 5 dpf showed significantly reduced locomotor activity and exploratory behaviour compared to the control (Fig. 2F, H, and I). In addition, VRAC knockdown larvae exhibited increased freezing behaviour (Fig. 2G) and thigmotaxis (Fig. 2J), consistent with elevated depression- and anxiety-like states.

### Transcriptional Consequences of VRAC Disruption in Zebrafish

RNA-seq analysis of *lrrc8aa* morphants and control zebrafish larvae (n = 3 per group) showed high and consistent alignment rates to the GRCz11 reference genome (>94.9% across all samples). Principal component analysis of normalized expression data revealed clear separation between groups along PC1, which accounted for 24.3% of the total variance (Fig. S1-A) Of 15,010 expressed genes, 4,282 (28.5%) were differentially expressed at a false discovery rate (FDR) < 0.05, including 2,078 upregulated and 2,204 downregulated genes (Fig. 3A). Within a curated set of 340 neurobiologically relevant genes (Listed in supplementary excel sheet), 21 satisfied stringent selection criteria (log₂ fold change ≥ 1 and FDR < 0.05). The most significantly upregulated gene was *rxfp3.3b* (log₂ fold change = 2.28; FDR = 1.27 × 10⁻⁸), a receptor associated with stress and arousal signaling. In contrast, *bhlhe22* exhibited the greatest downregulation (log₂ fold change = −2.34; FDR = 1.64 × 10⁻⁶), consistent with its role in cortical neuronal differentiation. Gene Ontology and KEGG pathway analyses indicated significant enrichment of terms related to synaptic signaling, regulation of ion transport, and cellular stress responses, suggesting coordinated pathway-level remodelling (Fig. 3B, Fig. S1-S2).

**Figure 3.**
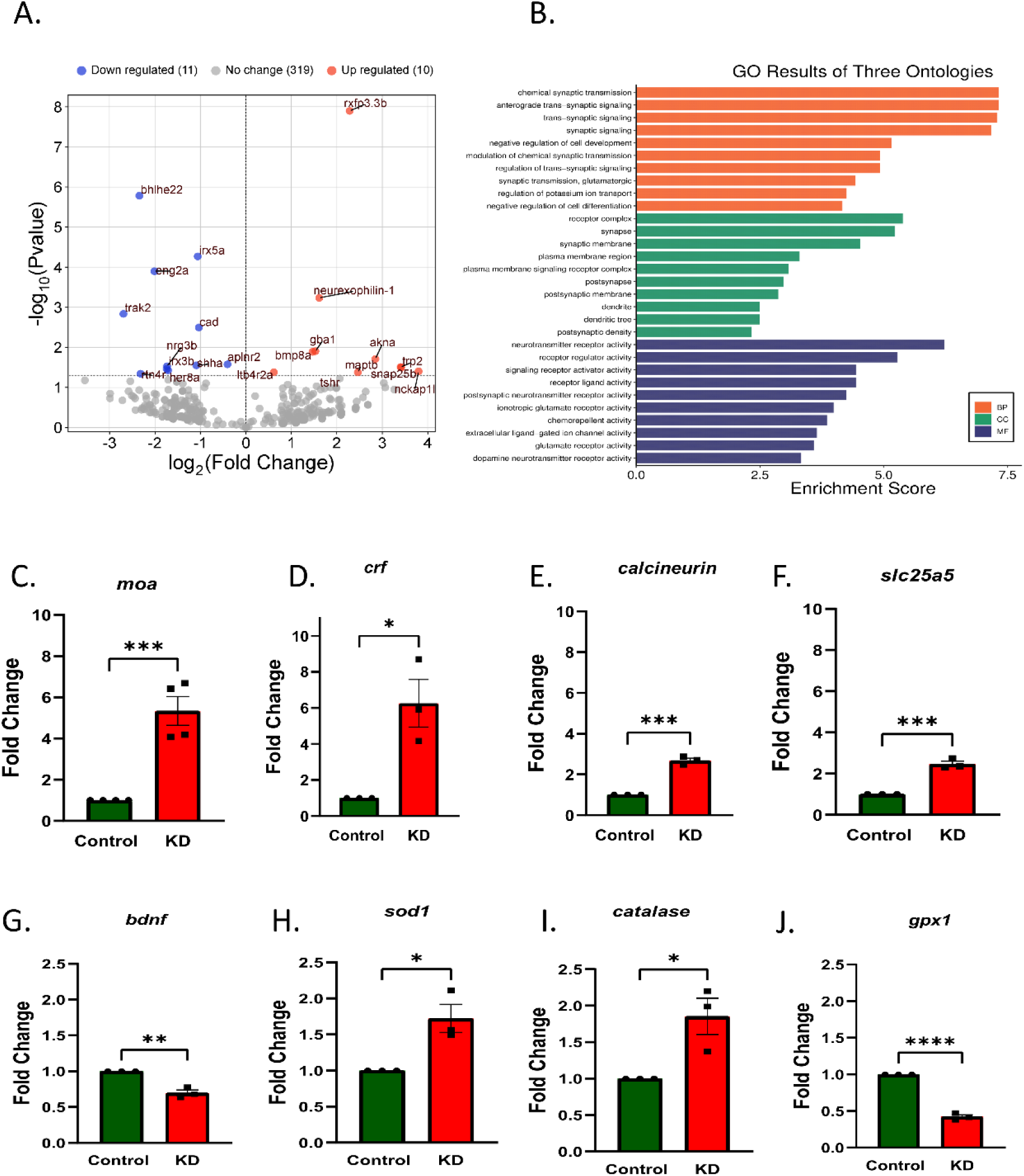
VRAC disruption alters the larval transcriptome and expression of depression - associated genes. (A)Volcano plot of differentially expressed genes in control and *lrrc8aa* knockdown larvae. Significantly upregulated (red) and downregulated (blue) genes (log₂FC ≥ 1, FDR < 0.05) are highlighted. (B) Gene Ontology enrichment analysis of deregulated genes. (C–G) qRT–PCR validation showing increased expression of *mao*, *crf*, calcineurin, and *slc25a5*, and decreased expression of *bdnf* in *lrrc8aa* knockdown (KD) larvae. BP, Biological Process; CC, Cellular Component; MF, Molecular Function. (H–J) Expression of oxidative stress-related genes showing increased *sod1* and *catalase*, and decreased *gpx1* expression. Data are presented as mean ± s.e.m. (N = 3–4 biological replicates; 50 pooled larval heads per replicate). Statistical significance was determined using an unpaired Student’s *t*-test.

Because transcriptome profiling may not reliably detect low-abundance transcripts, selected neuropsychiatric-relevant genes were validated using quantitative RT-PCR. *lrrc8aa*-depleted larvae showed significant upregulation of *mao*, *crf*, *calcineurin*, *slc25a5*, *sod1*, and *catalase*, together with downregulation of *bdnf* and *gpx1 (*Fig. 3C-J). This expression profile is consistent with molecular features associated with depressive-like states, including altered monoaminergic signaling, activation of the hypothalamic–pituitary–adrenal axis, reduced neurotrophic support and increased oxidative stress (Chakravarty et al., 2013; Karege et al., 2002; Sanders and Nemeroff, 2016).

### VRAC activator Zinc pyrithione (ZPT) rescues behavioural and molecular deficits in *lrrc8aa* knockdown larvae

To determine whether restoring VRAC activity could rescue the phenotypes observed in *lrrc8aa* knockdown larvae, we treated the larvae with ZPT, a potent VRAC activator (Figueroa and Denton, 2021). Larvae were treated with 5, 10, or 15 µM ZPT for 1 hour, followed by a 1-hour locomotor activity recording. ZPT improved locomotor activity in *lrrc8aa* knockdown larvae at all tested concentrations (Fig S3.A-C). As shown in Figures 4A–C, total distance travelled and high-velocity movement (≥8 mm/s, predefined high-activity threshold in the ViewPoint ZebraLab software.) were significantly increased in ZPT-treated knockdown larvae compared to untreated knockdown controls. In contrast, ZPT had minimal effect on locomotor behaviour in larvae treated with control morpholino. At the molecular level, the elevated *mao* mRNA expression observed in *lrrc8aa* knockdown brains was restored to control levels following ZPT treatment (Fig. 4D), indicating reversal of depression-associated transcriptional changes. Collectively, these findings demonstrate that pharmacological activation of VRAC can compensate for reduced channel activity and restore both behavioural and molecular phenotypes to baseline.

**Figure 4.**
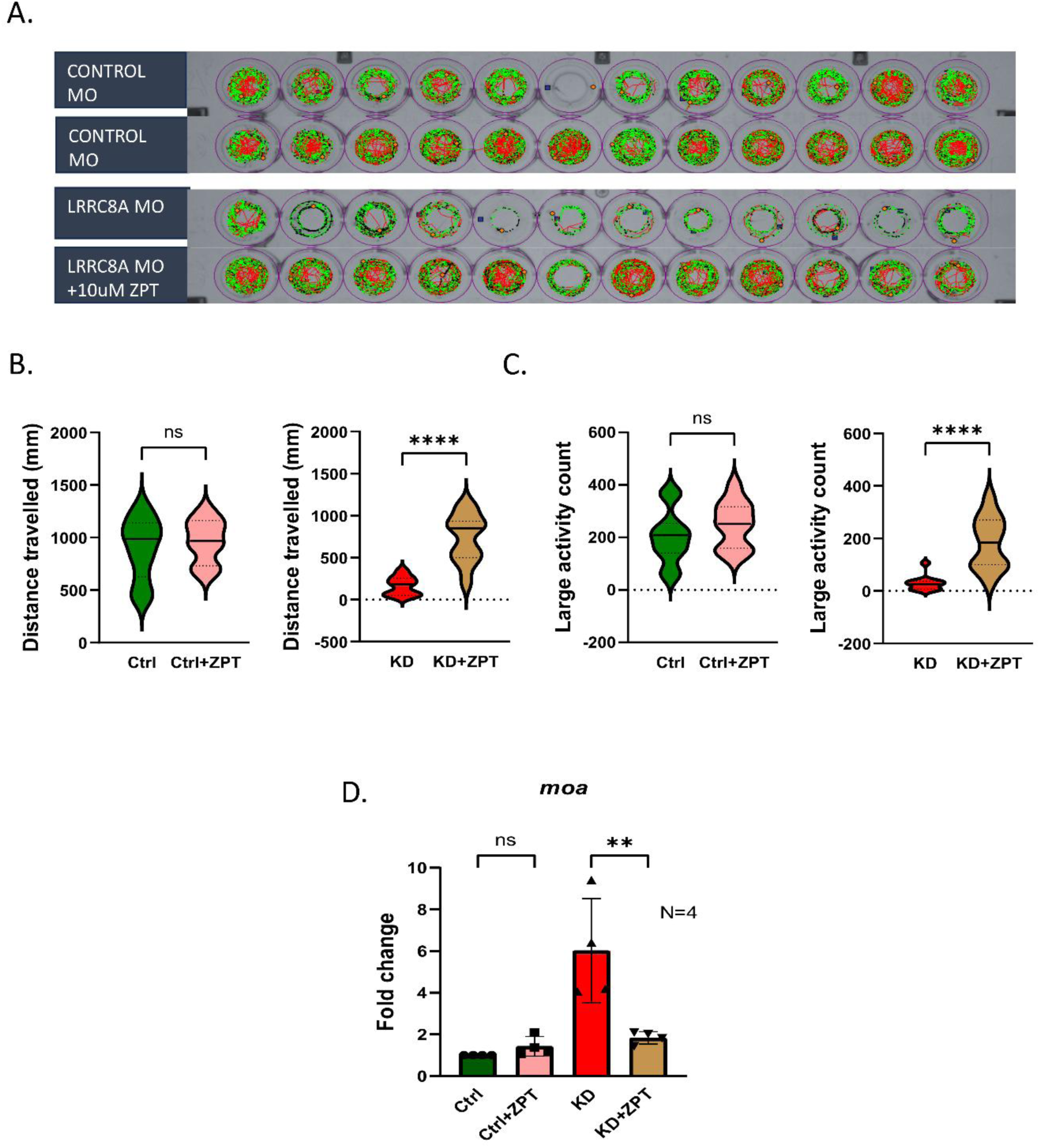
Pharmacological activation of VRAC rescues behavioural deficits and normalizes *mao* expression in *lrrc8aa*-depleted larvae. (A) Representative locomotor trajectories of control morpholino (MO)- and *lrrc8aa* MO-injected larvae treated with vehicle or ZPT (10 μM). (B–C) Quantification of total distance travelled and high-velocity movement events (≥8 mm/s), showing behavioural recovery in knockdown larvae following ZPT treatment (n = 11–12 larvae/group). (D) qRT–PCR analysis of *mao* expression, elevated in *lrrc8aa* knockdown larvae and restored by ZPT treatment. Data are presented as mean ± s.e.m. (N = 4 biological replicates; 50 pooled larval heads per replicate). Statistical analyses are described in Methods. C MO, control morpholino. Ctrl, control; KD, knockdown.

### VRAC activation alleviates depressive-like behaviour, restores neuronal morphology, and reduces elevated MAO levels in adult zebrafish

To induce depression-like behaviour in adult zebrafish, we employed a chronic unpredictable stress (CUS) paradigm and evaluated the effects of ZPT. CUS is a well-established model that reliably produces despair-like behaviour, anxiety, and cognitive deficits closely mirroring the core symptoms of human major depressive disorder (Chakravarty et al., 2013; Golla et al., 2020).

Anxiety- and depressive behaviours were assessed using the novel tank test (NTT), which measures freezing duration, zone preference, transitions, and swimming pattern. In this test, anxiety and depression are reflected by reduced exploration, increased freezing, erratic swimming, and prolonged bottom-dwelling. Control fish exhibited broad exploratory trajectories spanning the entire tank, whereas CUS-exposed fish remained predominantly in the lower half (Fig. 5A-B). Treatment with the reference antidepressant imipramine (IMP) restored normal exploration, and ZPT produced a comparable effect, with fish resuming full-tank exploration. Quantitative analysis confirmed these observations (Fig. 5C-E). CUS-induced increases in freezing time, reduced upper-zone occupancy, and increased transitions to the lower zone were all reversed by ZPT to an extent comparable to IMP (Fig. 5C-E) Additionally, CUS-elevated meandering frequency, which reflects erratic anxiety-driven swimming, was significantly reduced by ZPT but not by IMP, though IMP showed a non-significant trend toward reduction (Fig. 5F). This suggests that ZPT may offer broader anxiolytic benefits than imipramine. Aggression was assessed using the mirror-induced aggression test, where aggression was defined as mirror-directed fast swimming and biting. CUS markedly reduced mirror-zone occupancy, indicating suppressed aggressive drive. ZPT partially restored this behaviour to a level comparable to IMP (Fig. 5G). In the light-dark preference test, CUS fish spent significantly less time in the light zone, consistent with heightened anxiety, and both ZPT and IMP partially reversed this preference ( Fig. 5H). Together, these findings demonstrate that ZPT exerts anxiolytic and antidepressant -like effects in the zebrafish CUS model.

**Figure 5.**
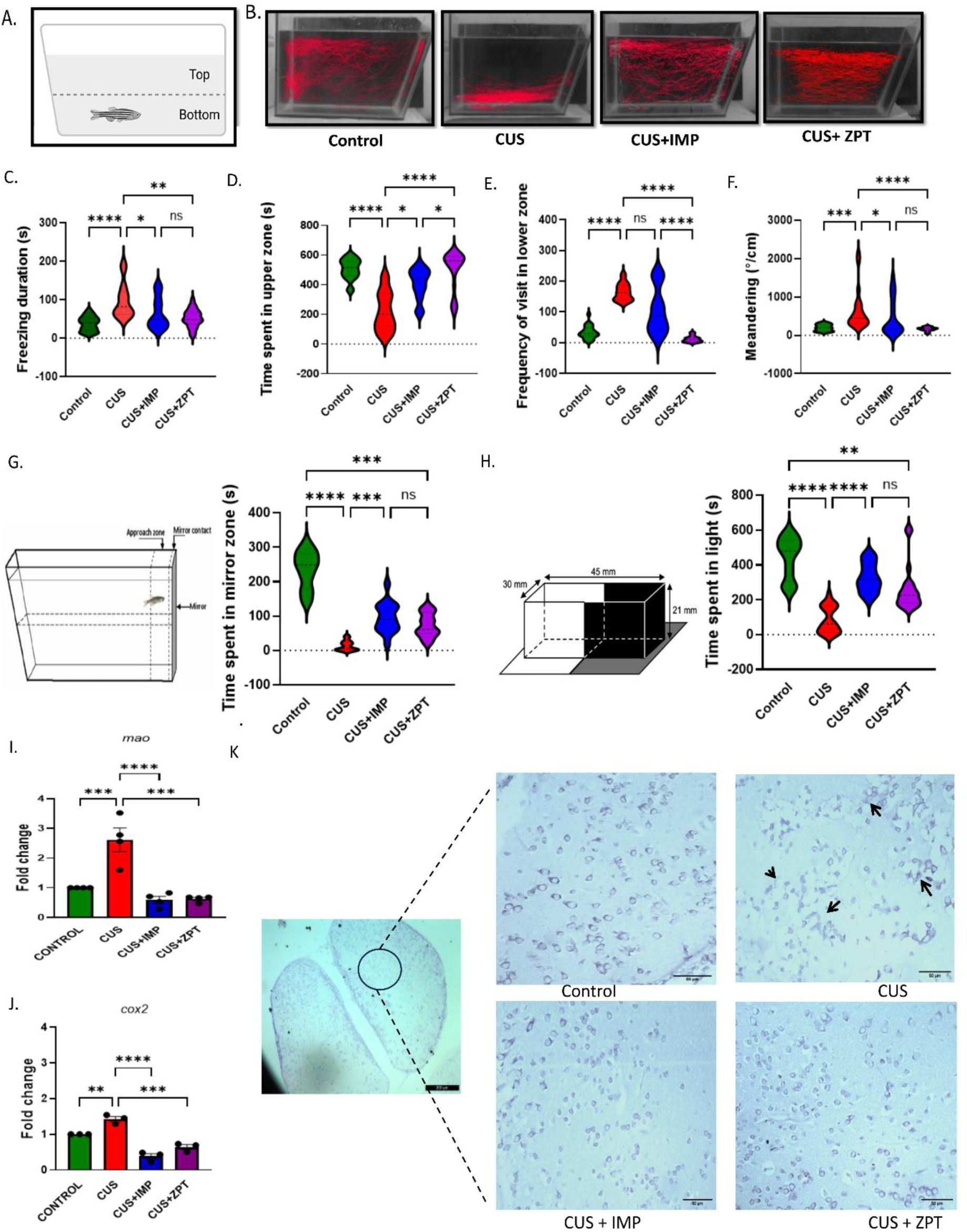
VRAC activation reverses chronic stress-induced behavioural, molecular, and neuronal alterations in adult zebrafish. (A) Schematic of the novel tank test (NTT). (B) Representative locomotor trajectories of control, chronic unpredictable stress (CUS), CUS + imipramine (IMP), and CUS + zinc pyrithione (ZPT) groups. (C–F) NTT behavioural measures: freezing duration, time in the upper zone, lower-zone entries, and meandering (n = 20 fish/group). (G) Mirror-induced aggression assay and (H) light–dark preference test (n = 15–20 fish/group). (I–J) Brain qRT–PCR analysis showing *mao* and *cox2* expression, elevated by CUS and restored by ZPT or imipramine (N = 3–4 biological replicates). (K) Representative cresyl violet-stained brain sections showing CUS-induced neuronal shrinkage and its reversal by ZPT and imipramine. Arrows indicate altered neurons. Scale bar, 50 μm. Statistical analyses are described in Methods.

To confirm that the observed behavioural effects were mediated by the intact Zn^2+^-pyrithione complex rather than free Zn^2+^, CUS-exposed fish were treated with ZnCl_2_. ZnCl_2_ did not improve any behavioural phenotype (Fig. S4), consistent with previous reports that free Zn^2+^ alone does not activate VRAC (Figueroa and Denton, 2021). Furthermore, ZPT or IMP treatment in non-stressed control fish did not alter baseline behavioural parameters, indicating that neither treatment disrupts normal behaviour under unstressed conditions.

We next investigated whether ZPT could normalize the CUS-induced changes in gene expression in brain tissue. CUS increased *mao* mRNA levels approximately 2.5-fold relative to control animals; treatment with either ZPT or IMP restored *mao* mRNA to control levels (Fig. 5I). Similarly, CUS produced a moderate increase in *cox2* mRNA expression, a marker of neuroinflammation, and this increase was also attenuated by both ZPT and IMP treatment (Fig. 5J). To assess neuronal integrity, brain sections were stained with cresyl violet. CUS fish showed reduced neuronal soma size, suggesting neuronal atrophy associated with chronic stress exposure (Fig. 5K). ZPT treatment markedly improved neuronal morphology, restoring cell size and structural integrity to levels comparable to IMP-treated fish (Fig.5K)

## Discussion

Our findings indicate that reduced VRAC function contributes to affective dysregulation in zebrafish. Both pharmacological inhibition and genetic depletion of VRAC produced similar behavioural abnormalities, including reduced locomotion, increased freezing, and enhanced thigmotaxis, together with molecular changes associated with depressive states. Importantly, restoration of VRAC activity reversed many of these behavioural and molecular alterations. A major strength of this study is the use of complementary pharmacological, genetic, electrophysiological, transcriptomic, and behavioural approaches. The similar phenotypes produced by DCPIB treatment and *lrrc8aa* knockdown strongly support a role for VRAC in affective regulation. Because touch-evoked escape responses remained intact, the observed behavioural changes are unlikely to reflect general motor impairment. Consistent with previous studies, freezing and thigmotaxis are more likely to represent anxiety- and depression-like states than locomotor deficits (Maximino et al., 2010).

VRAC regulates cell volume and mediates the release of neuroactive molecules, including taurine and glutamate. Therefore, reduced VRAC activity could potentially disrupt neurotransmitter homeostasis and contribute to the behavioural abnormalities observed in this study. Transcriptomic analysis further revealed alterations in pathways related to synaptic signalling, ion transport, and cellular stress responses. Changes in *mao*, *crf*, *bdnf*, and *gpx1* expression are consistent with molecular pathways frequently implicated in depression, including altered monoaminergic signalling, stress-axis activation, reduced neurotrophic support, and oxidative stress. The increase in *sod1* and catalase together with reduced *gpx1* suggests activation of a compensatory antioxidant response following VRAC deficiency. The ability of ZPT to restore VRAC currents, improve behaviour, and normalize elevated *mao* expression further supports a functional link between VRAC activity and affective state. Importantly, ZPT had minimal effects in control larvae, arguing against a non-specific stimulatory action.

The adult chronic unpredictable stress model extends these findings beyond larval development. ZPT improved multiple behavioural measures to an extent comparable to imipramine and also reduced stress-induced meandering behaviour, an effect that was less evident with imipramine. This observation suggests that VRAC activation and monoaminergic antidepressants may act through partially distinct mechanisms. In addition, ZPT improved stress-induced neuronal morphological abnormalities, consistent with a role for VRAC in maintaining cellular homeostasis under chronic stress conditions.

Together, our findings identify VRAC as a potential non-monoaminergic target for mood disorders. Unlike conventional antidepressants, which primarily modulate monoamine signalling, VRAC may influence affective behaviour through regulation of cellular homeostasis, glial-neuronal communication, and neurotransmitter release. Given the limitations of current antidepressant therapies, further investigation of VRAC as a therapeutic target appears warranted. Future studies should determine whether alterations in LRRC8 expression or genetic variation are associated with depression and anxiety disorders in humans.

## Limitations

Several limitations should be considered when interpreting these findings. First, morpholino-mediated knockdown is transient and may produce developmental effects; future studies should employ stable CRISPR-generated *lrrc8aa* loss-of-function zebrafish lines. Second, our experiments do not distinguish between the contributions of neuronal and astrocytic VRAC, which will require cell-type-specific genetic approaches. Third, although the ZnCl₂ control and electrophysiological data support VRAC as a major target of ZPT, additional pharmacological actions of ZPT cannot be completely excluded. Furthermore, behavioural assays in zebrafish provide indirect measures of affective state, and extrapolation to human depression should therefore be made cautiously. Finally, validation of these findings in mammalian models of depression will be necessary to establish their broader translational relevance.

## Supporting information

List of selected DEG-transcriptome data

Supplementary video 1 Touch Response in DCPIB treated Larvae

Supplementary video 2 Touch Response in lrrc8aa KD Larvae

## Acknowledgements

We acknowledge Kalaiarasan and Amit Kumar Ghosh for their critical comments on the manuscript.

## Author contribution statement

AA conceptualized experiments, acquired data, analysed data, interpreted data, and wrote the manuscript. DS acquired data, analysed data, interpreted data, and wrote the manuscript. RS acquired and analysed data. GM acquired data. AKB conceptualized experiments, provided funding, interpreted data, and wrote the manuscript.

## Declaration of generative AI and AI-assisted technologies in the writing process

During the preparation of this work, the authors used ChatGPT to improve readability and language. After using this tool, the authors reviewed and edited the content as needed and take full responsibility for the content of the publication.

## Data availability statement

The data used to support the findings of this study are available from the corresponding author upon request.

