## Supplementary Data for "Altered volume-regulated anion channel activity contributes to depression and anxiety-related molecular and behavioural phenotypes in zebrafish"

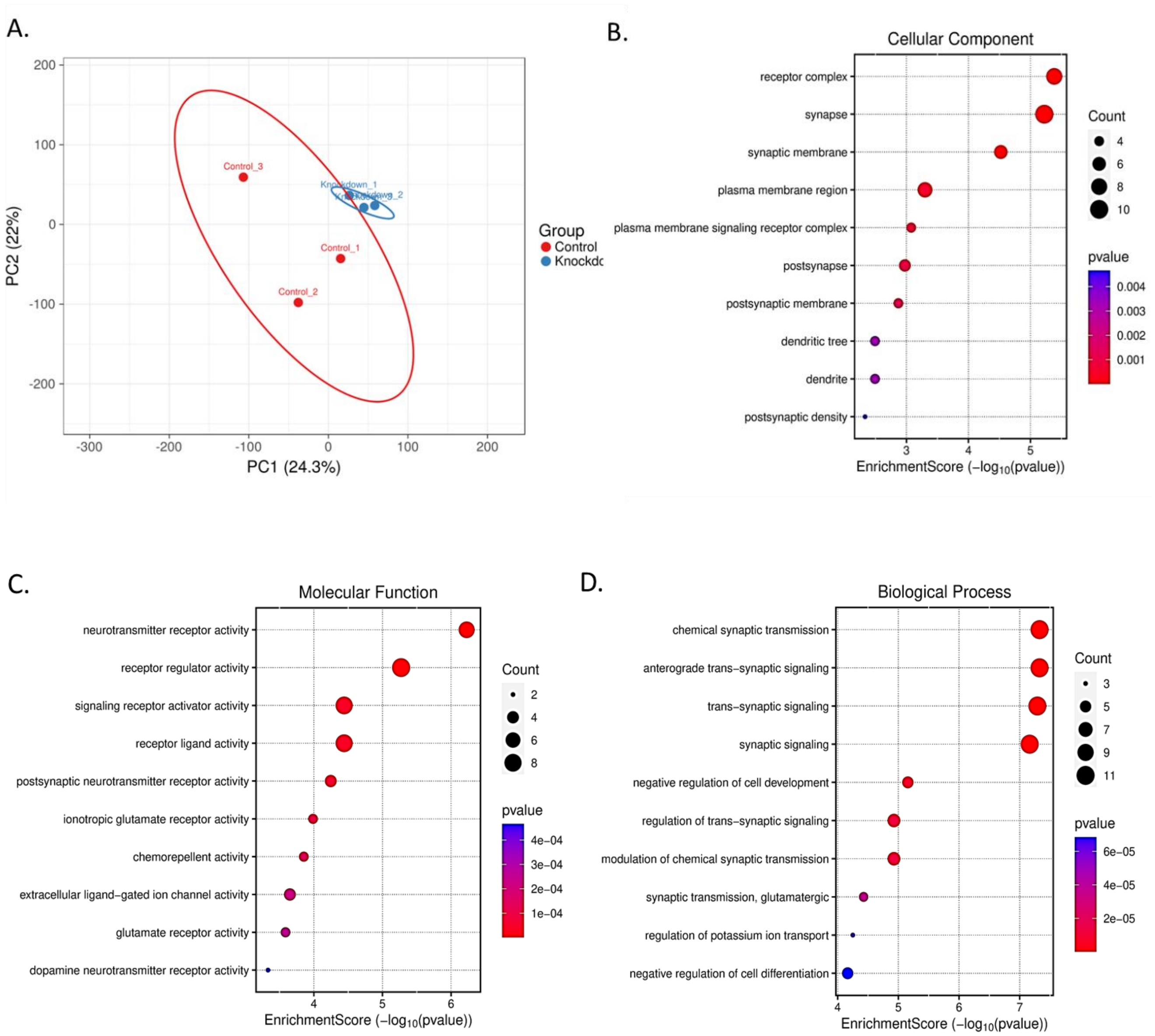


**Figure S1. Transcriptomic profiling of control and *lrrc8aa* knockdown larvae** (A) PCA plot of normalized RNA-seq data showing distinct clustering of control and *lrrc8aa* knockdown samples (n = 3 per group). PC1 and PC2 explain 24.3% and 22% of the variance, respectively. (B–D) Dot plots of enriched Gene Ontology terms for Biological Process, Cellular Components, and Molecular Function. Dot size indicates gene count and colour denotes statistical significance.

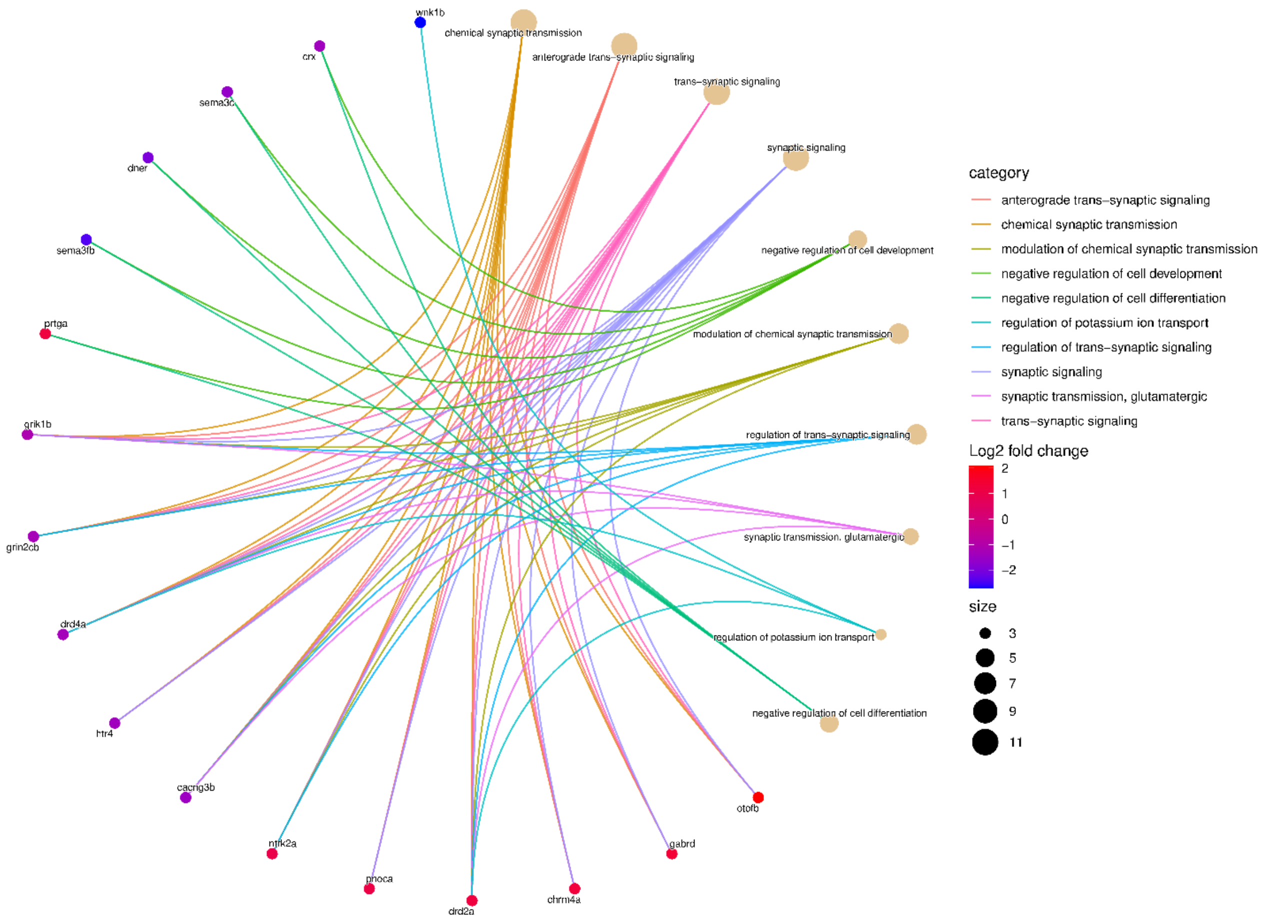


**Figure S2. Gene–pathway network of differentially expressed genes and enriched biological processes**

Nodes represent genes and Gene Ontology terms, with edges indicating gene–pathway associations. Edge colour denotes the direction and magnitude of gene expression changes ($log_2FC$).

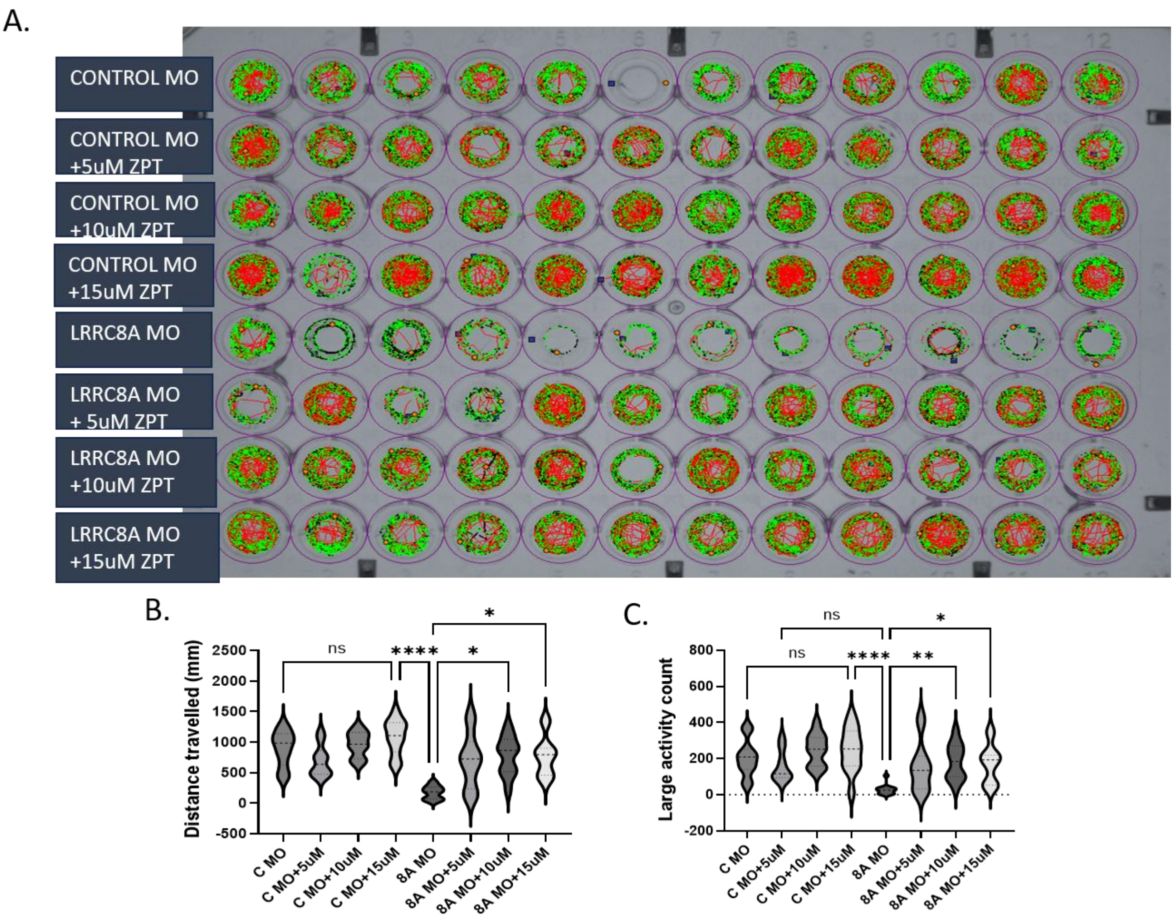


**Figure S3. Dose-dependent effects of zinc pyrithione (ZPT) on locomotor activity in zebrafish larvae**

(A) Representative locomotor trajectories from the 96-well open-field assay for control morpholino (C MO)- and *lrrc8aa* morpholino (8A MO)-injected larvae treated with ZPT (5, 10, or 15 µM). Green and red tracks indicate normal- and high-activity movements, respectively. (B) Total distance travelled and (C) high-activity events per larva across experimental groups (n = 12 larvae/group). Data were analyzed using the Kruskal–Wallis test followed by Dunn’s multiple comparisons test.

.

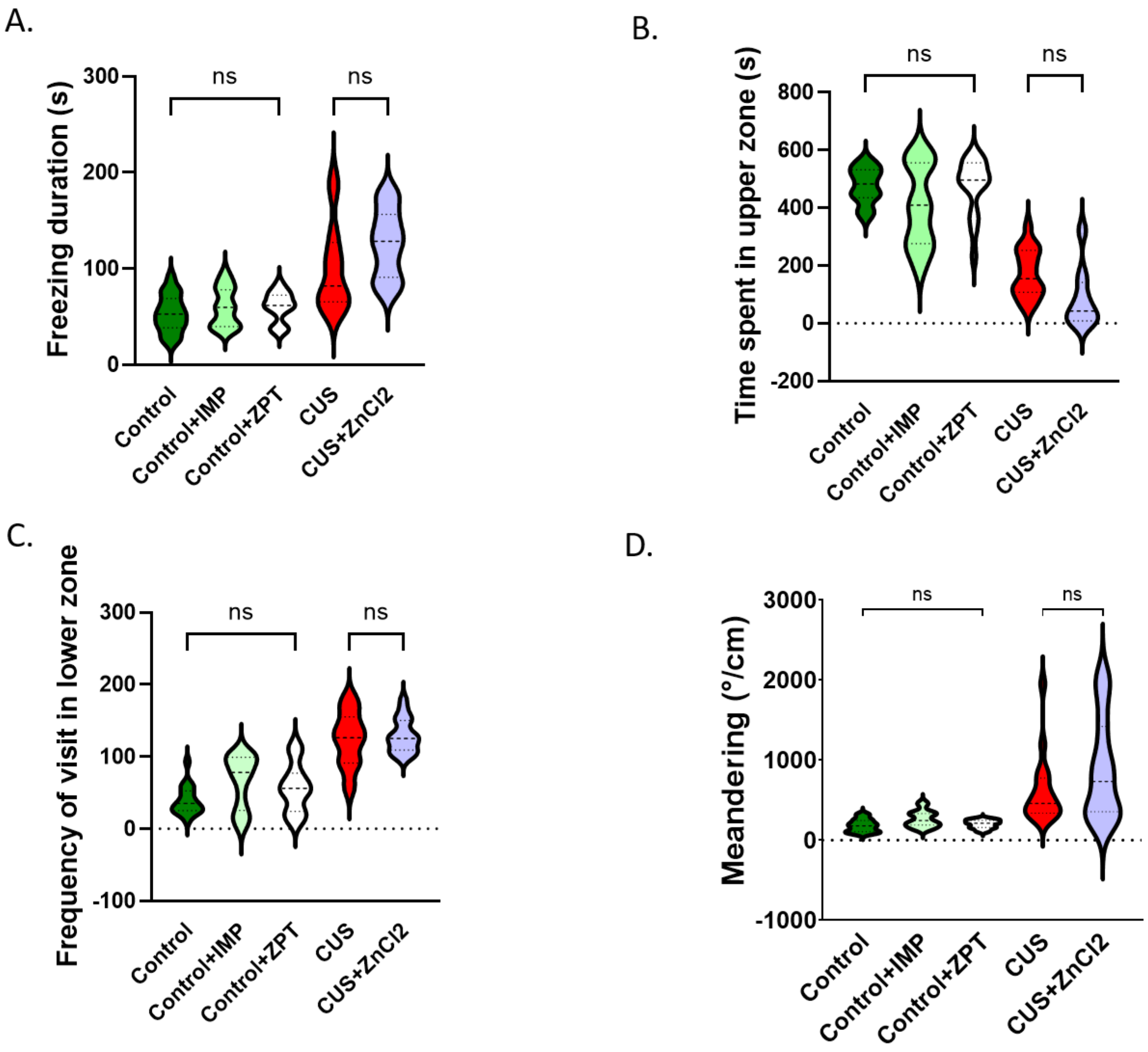


**Figure S4. Effects of $ZnCl_2$, ZPT, and imipramine (IMP) in the novel tank test (NTT)** Violin plots showing (A) freezing duration, (B) time spent in the upper zone, (C) frequency of entries into the lower zone, and (D) meandering behaviour in adult zebrafish (n = 17 fish per group). Data are presented as mean ± s.e.m. Statistical significance was assessed using the Kruskal–Wallis test followed by Dunn's multiple comparisons test due to non-normal distribution in at least one group. CUS, Chronic unpredictable stress.

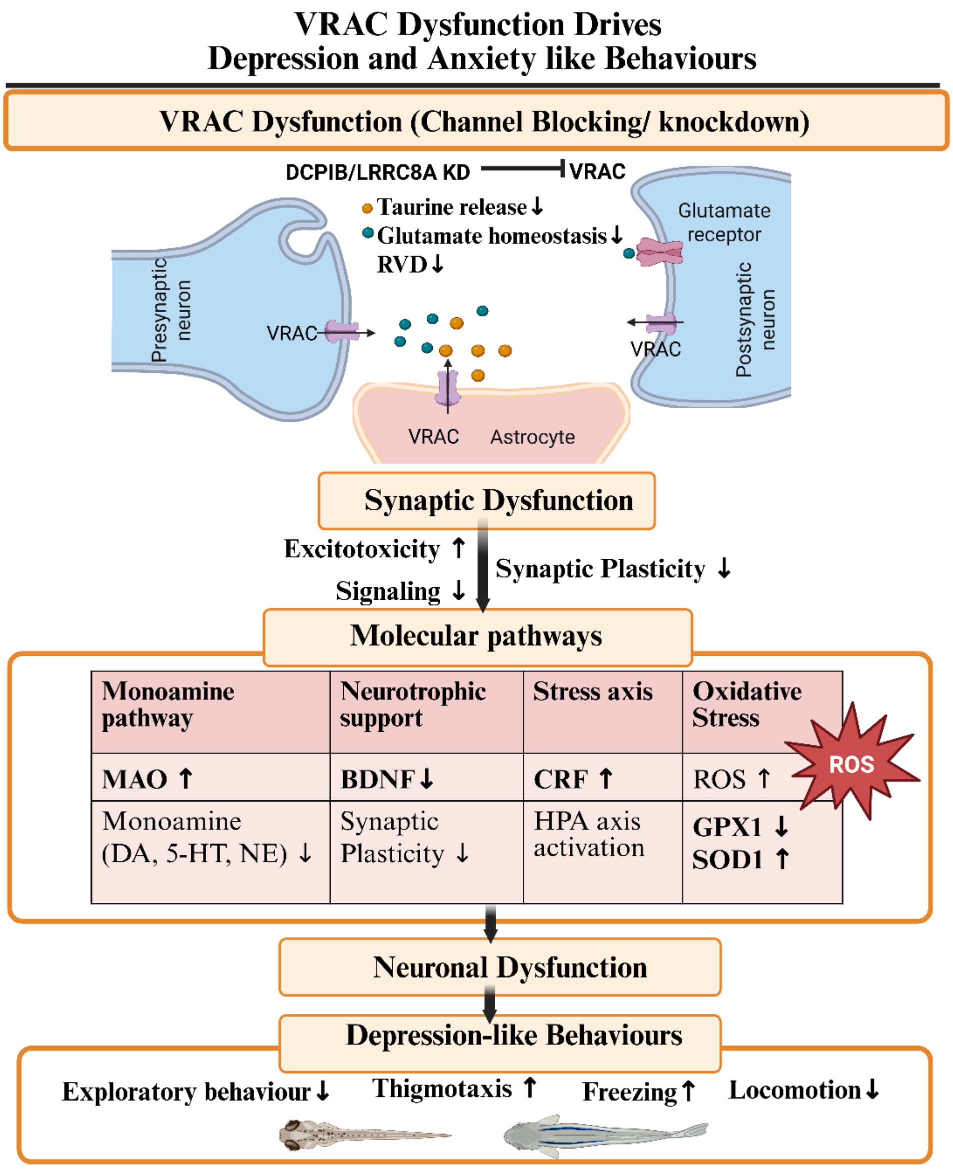


**Figure S5. Proposed Mechanism Linking Reduced VRAC Function to Affective Behavioural Alterations in Zebrafish** (Created with BioRender).

**Table 1**

**CUS paradigm for developing anxiety/ depression like phenotype in zebrafish**

| | **Day 1** | **Day 2** | **Day 3** | **Day 4** | **Day 5** | **Day 6** | **Day 7** | **Day 8** | **Day 9** | **Day 10** | **Day 11** | **Day 12** | **Day 13** | **Day 14** | **Day 15** |
|---|---|---|---|---|---|---|---|---|---|---|---|---|---|---|---|
| **Morning** | RS | SI | OC | NE | CS | CN | DBE | HS | WC | SI | RS | OC | OC | WC | SI |
| **Evening** | TC | HS | CN | DBE | RS | WC | TC | OC | NE | CS | DBE | TC | CN | NE | HS |
| | | | | | DT | DT | DT | DT | DT | DT | DT | DT | DT | DT | DT |

RS – Restraint Stress; SI – Social Isolation; OC – Overcrowding; TC – Tank change; CS – Cold stress; CN – Chasing with Net; HS – Heat stress; DBE – Dorsal Body Exposure; WC- Water Change, NE – net elevation, DT-Drug Treatment.

**Table 2**

**List of primers used**

| Target gene | Forward Primer (5′>3′) | Reverse Primer (5′>3′) |
|---|---|---|
| *lrrc8aa* | GAGCGGGATTCCTGATACGG | GCTGAATATGGAGTGCGGGA |
| *mao a* | GTGTGCAATGGGCAAACTCC | GAATCCTCCAACCGATGGCA |
| *bdnf* | GGACAAAAAGACGGCAATAGAC | CGATCTTCCTTTTGCTATCCAT |
| *crf* | CATCCCAGTATCCAAAAAGAGC | TCGTAGCAGATGAAAGGTCAGA |
| *calcineurin* | GCCTTTAGGATCTACGACATGG | ATATTCTCCCGTCTCCGTCTTT |
| *slc25a5* | CATCATTTACAGAGCTGCCTAC | TTTACGTCCAGACTGCATCATC |
| *sod1* | ACCGGCACCGTCTATTTCAA | AGCATGGACGTGGAAACCAT |
| *catalase* | AAAATGGGGGCCTTTGCATAC | GCAGAAAGGACGGCAAACATT |
| *gpx1a* | TTTACGACCTGTCCGCGAAA | CTGTTGTGCCTCAAAGCGAC |
| *beta-actin* | CCTTCCAGCAGATGTGGATTAG | TGAAGTGGTAACAGTCCGTTTAG |
| *cox2* | GCGGCCGAATTCAATACCCT | GTTACGTCCACCAGACACCC |
